# Temporal cell fate determination in the spinal cord is mediated by the duration of Notch signalling

**DOI:** 10.1101/2021.10.25.465818

**Authors:** Craig T. Jacobs, Aarti Kejriwal, Katrinka M. Kocha, Kevin Y. Jin, Peng Huang

## Abstract

During neural development, progenitor cells generate different types of neurons in specific time windows. Despite the characterisation of many of the transcription factor networks involved in these differentiation events, the mechanism behind their temporal regulation is poorly understood. To address this question, we studied the temporal differentiation of the simple lateral floor plate (LFP) domain in the zebrafish spinal cord. LFP progenitors sequentially generate early-born Kolmer-Agduhr” (KA”) interneurons and late-born V3 interneurons. Analysis using a Notch signalling reporter demonstrates that these cell populations have distinct Notch signalling profiles. Not only do V3 cells receive higher total levels of Notch response, but they collect this response over a longer duration compared to V3 cells. To test whether the duration of Notch signalling determines the temporal cell fate specification, we combined a transgene that constitutively activates Notch signalling in the ventral spinal cord with a heat shock inducible Notch signalling terminator to switch off Notch response at any given time. Sustained Notch signalling results in expanded LFP progenitors while KA” and V3 interneurons fail to specify. Early termination of Notch signalling leads to exclusively KA” cell fate, despite the high level of Notch signalling, whereas late attenuation of Notch signalling drives only V3 cell fate. This suggests that the duration of Notch signalling is instructive in cell fate specification. Interestingly, knockdown experiments reveal a role for the Notch ligand Jag2b in maintaining LFP progenitors and limiting their differentiation into KA” and V3 cells. Our results indicate that Notch signalling is required for neural progenitor maintenance while a specific attenuation timetable defines the fate of the postmitotic progeny.

## INTRODUCTION

Throughout spinal cord development, neural progenitors differentiate into their post-mitotic progeny in a strict spatiotemporal manner. Patterning of neural progenitor domains along the dorsal-ventral (DV) axis is achieved by combined actions of three major cell signalling pathways, Bone Morphogenic Protein (BMP), Wnt and Hedgehog (Hh) (Le Dréau and Martí, 2012). A gradient of BMP/Wnt from the roof plate patterns the dorsal spinal cord, while a gradient of Hh originating from the notochord and the floor plate is required for ventral fate specification (Andrews et al., 2019; Dessaud et al., 2008; Le Dréau and Martí, 2012; Ulloa and Martí, 2010). Neural progenitor domains can be identified by their expression of specific, highly conserved, transcription factors which in turn form a regulatory network that maintains the DV pattern (Briscoe and Small, 2015; Cohen et al., 2013; Jessell, 2000). Importantly, common progenitors in a single domain can give rise to different neuronal subtypes over time. However, how temporal cell fate specification is regulated is not well understood.

An essential mechanism for regulating neural cell diversity is mediated by Notch signalling through direct cell-cell communications (Louvi and Artavanis-Tsakonas, 2006; Pierfelice et al., 2011). The Delta and Jagged/Serrate family of ligands, present at the membrane of the “sending” cell, activate the Notch receptor at the membrane of the “receiving” cell, leading to the cleavage of the Notch intracellular domain (NICD). Once released, the NICD translocates to the nucleus where it complexes with the DNA binding protein RBPJ and the Mastermind-like (MAML) coactivator to form a ternary transcription activation complex that drives expression of downstream targets, such as the Hes/Hey family of transcription factors (Artavanis-Tsakonas and Simpson, 1991; Kopan and Ilagan, 2009; Pierfelice et al., 2011).

The canonical role for Notch signalling during neural development is to maintain progenitor state through preventing the expression of proneural genes (Guruharsha et al., 2012; Louvi and Artavanis-Tsakonas, 2006; Yoon and Gaiano, 2005). This role is highly conserved and has been well classified in organisms ranging from *Drosophila* to mouse (Formosa-Jordan et al., 2013; Kageyama et al., 2008). For example, in early nervous system development in *Drosophila*, Notch-mediated lateral inhibition prevents the expression of the proneural transcription factor achaete-scute complex (AS-C) leading to a “salt and pepper” pattern of neural precursors and neuroepithelial cells (Sato et al., 2016). In vertebrates, the blockade of Notch signalling through multiple methods results in the progressive loss of progenitors, while the constitutive activation of Notch signalling inhibits the formation of differentiated neuronal populations (Appel et al., 2001; Huang et al., 2012; Park and Appel, 2003; Yang et al., 2006). In addition to its role in maintaining neural progenitors, Notch signalling has also been implicated in binary neuronal fate decisions. For example, active Notch signalling promotes the inhibitory V2b interneuron fate over the excitatory V2a fate from the same progenitors (Batista et al., 2008; Del Barrio et al., 2007; Kimura et al., 2008; Okigawa et al., 2014; Peng et al., 2007). Similarly, motor neuron precursors (pMN) with higher level of Notch signalling differentiate into Kolmer-Agduhr’ (KA’) interneurons, whereas those with lower Notch activity develop as primary motor neurons (Shin et al., 2007). These studies suggest that the level of Notch signalling contributes to the cell type diversification of the spinal cord. More recently, we and others have shown that Notch signalling also functions to maintain the Hh responsiveness of neural progenitors (Huang et al., 2012; Jacobs and Huang, 2019; Kong et al., 2015; Stasiulewicz et al., 2015). PHRESH (PHotoconvertible REporter of Signalling History) analysis reveals stereotypic temporal dynamics of Notch signalling during spinal cord development (Jacobs and Huang, 2019). Indeed, the attenuation of both Notch and Hh signalling is required for the proper differentiation of lateral floor plate (LFP) progenitors into Kolmer-Agduhr” (KA”) interneurons (Huang et al., 2012). This raises the question whether the duration of Notch signalling also plays a role in cell fate determination in the spinal cord.

In mouse and chick, Hh signalling induces the expression of the transcription factor *Nkx2.2* in the p3 domain, immediately dorsal to the floor plate (Briscoe et al., 2000). During patterning, this domain contains V3 progenitors medially, while terminally differentiated V3 interneurons reside laterally (Alaynick et al., 2011; Dessaud et al., 2008). In the zebrafish, the analogous, single cell-width lateral floor plate (LFP) domain flanks the medial floor plate (MFP) on either side of the midline (Odenthal et al., 2000). Similar to the p3 domain of the chick and mouse, the LFP domain contains both progenitors and differentiated interneurons (Lewis and Eisen, 2003; Schäfer et al., 2005; Schäfer et al., 2007). Interestingly, there are two distinct types of interneurons present: early born KA” interneurons and later born V3 interneurons. Previously thought to be zebrafish specific, cells analogous to KA” interneurons have recently been described as cerebrospinal fluid contacting neurons (CSF-cNs) in mouse (Petracca et al., 2016). The transcription factors and genetic markers that describe KA” cell fate have been well established (Yang et al., 2010; Yang et al., 2020), while the genetic programs defining V3 interneurons in zebrafish are not well understood. Moreover, although V3 cells are located in the LFP domain (Schäfer et al., 2007), their lineage relationships with LFP progenitors and KA” cells remain unclear.

Here, we describe the differentiation dynamics of V3 interneurons during LFP patterning in zebrafish and demonstrate their lineage relationship to KA” interneurons. Using our previously described Notch signalling reporter (Jacobs and Huang, 2019), we show that the three cell types in the LFP domain have distinct duration of Notch signalling. To analyse the importance of duration of response in comparison to total level, we develop a specific transgenic tool that allows us to artificially sustain Notch signalling in the ventral spinal cord before conditionally attenuating it. This Notch^ON^-Notch^OFF^ method demonstrates that to acquire V3 fate over KA” fate, LFP progenitors must sustain Notch signalling for a prolonged duration. Interestingly, knockdown of *jag2b* results in expansion of both KA” and V3 cells within their normal differentiation windows. Together, our data reveal that Notch signalling functions to provide a timetable that guides fate decisions during LFP patterning.

## RESULTS

### Differentiation dynamics of V3 interneurons in the LFP domain

The lateral floor plate (LFP) is a single cell wide domain that flanks the medial floor plate at the ventral extent of the zebrafish spinal cord. It begins as a homogenous population of LFP progenitors, but by 48 hpf it consists of two populations of differentiated neurons, KA” and V3 interneurons, alongside LFP progenitors (Fig. 1A). We combined two transgenic reporters in order to visualise the post-mitotic neuronal populations – *gata2:GFP* for KA” and *vglut2a:dsRed* for V3 interneurons – during LFP patterning (Fig. 1B). At 24 hpf, only KA” cells were present in the LFP domain, with V3 cells arising at the lateral edge of the domain beginning at 30 hpf. Between 30 hpf and 48 hpf, the V3 population continued to expand and began to occupy the more medial space, while the number of KA” cells did not increase (Fig. 1B). Interestingly, when we used whole-mount in situ hybridisation for *sim1*, another marker for V3 cells, we could visualise V3 cells at the anterior extent of the spinal cord beginning at 24 hpf (Fig. 1C). This population slowly expanded along the anterior-posterior (AP) axis until reaching maximal coverage by 48 hpf (Fig. 1D). This is in stark contrast to KA” differentiation, which occurred rapidly along the AP axis between 16 hpf and 24 hpf, and reached a plateau after 24 hpf (Fig. 1E). This result suggests that KA” differentiation precedes the generation of V3 cells in the LFP domain.

**Figure 1.**
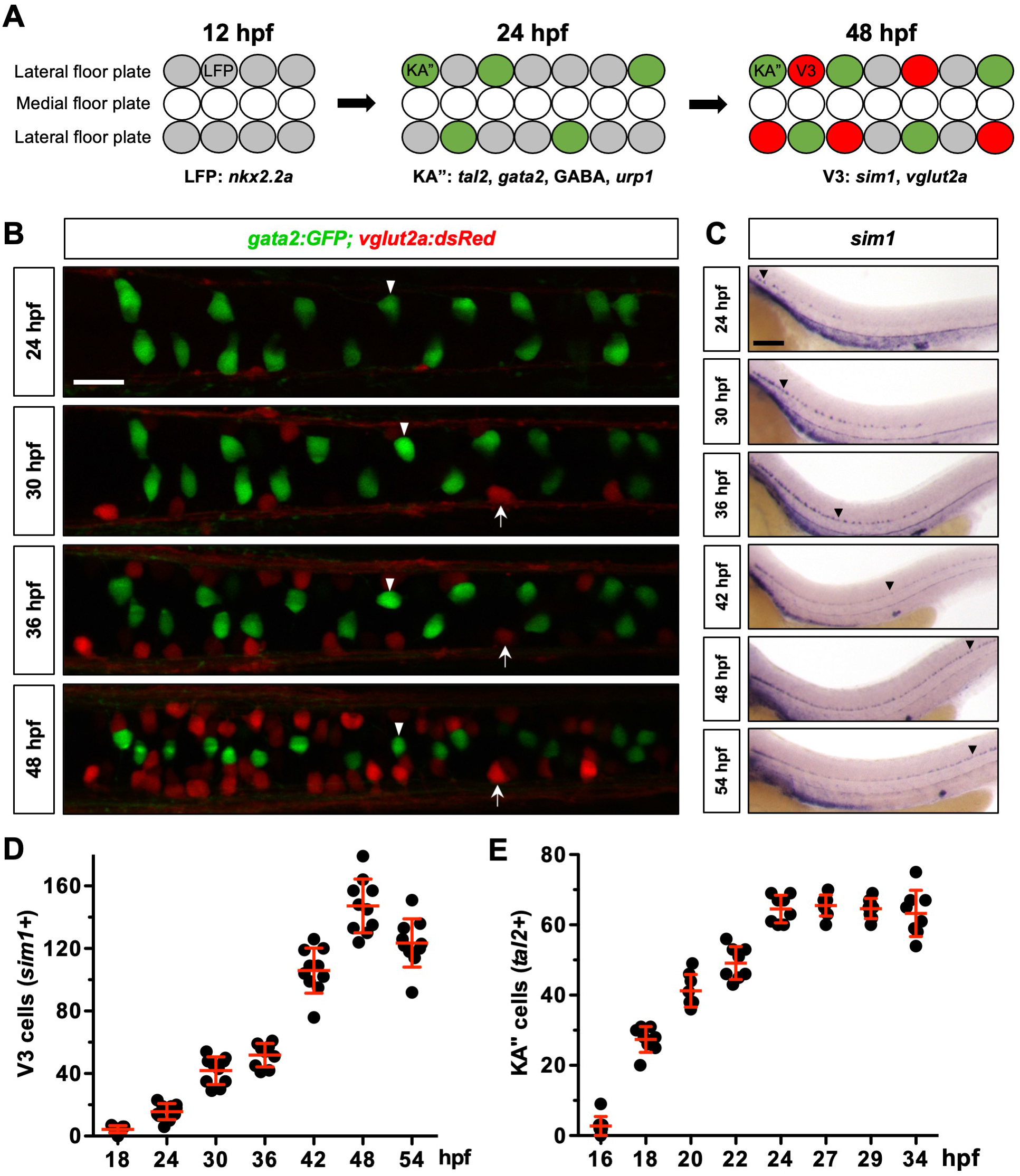
Characterisation of V3 and KA” interneurons in the LFP domain. (A) Schematic representation of a dorsal view of the ventral spinal cord at different stages during neuronal differentiation. The lateral floor plate domain, eventually consisting of LFP progenitors, KA” and V3 interneurons, flanks the medial floor plate at the ventral extent of the spinal cord. Specific markers for LFP, KA” and V3 cells are indicated. (B) *gata2:GFP; vglut2a:dsRed* embryos were followed and imaged from a dorsal view at 24, 30, 36, and 48 hpf. *gata2:GFP* and *vglut2a:dsRed* marks KA” (arrowheads) and V3 (arrows) interneurons, respectively. *n* = 3 embryos. (C) Whole-mount in situ hybridisation for *sim1* was performed on wild type embryos fixed at 6-hour intervals from 18 to 54 hpf. The timepoints between 24 and 54 hpf are shown. Arrowheads denote the extent of continuous V3 coverage. (D) The number of *sim1^+^* V3 cells were counted at each timepoint from the experiment described in (C). Each data point represents a single embryo. *n* = 10 embryos per timepoint. (E) The number of *tal2^+^* KA” cells were counted between 16 hpf and 34 hpf. Each data point represents a single embryo. *n* = 6-8 embryos per timepoint. Data are plotted with mean ± SD. Scale bars: (B) 20 μm; (C) 100 μm.

To examine the differences between the *vglut2a:dsRed* reporter and *sim1* as a V3 marker, we analysed *sim1* expression alongside the transgenic reporter in the anterior portion of the spinal cord (Fig. S1A). At 36 hpf, 33% of *sim1^+^* V3 cells were *vglut2a:dsRed^+^*, but by 48 hpf this proportion had risen to 91% (Fig. S1D). Similarly, only 40% of the *sim1^+^* population at 30 hpf was positive for the differentiated neuronal marker HuC, but the percentage increased to 60% by 48 hpf (Fig. S1B, D). Strikingly, at 30 hpf 75% of the *sim1^+^* population remained positive for the LFP progenitor marker *nkx2.2a*. At 48 hpf, this had significantly decreased, with only 17% of the *sim1^+^* V3 cells remaining *nkx2.2a^+^* (Fig. S1C, D). Taken together, this suggests that *sim1* is an early marker, while *vglut2a:dsRed* is a late marker, for V3 cell fate.

We have previously shown that KA” cells are derived from LFP progenitors (Huang et al, 2012). Considering that at 30 hpf about 75% of V3 cells also expressed the progenitor marker *nkx2.2a*, we hypothesised that V3 cells are derived from LFP progenitors that did not differentiate into KA” cells. To further interrogate this, we inhibited Notch signalling using the small molecule LY-411575 (Fauq et al., 2007). Notch inhibition causes premature differentiation and depletes the progenitor pool (Jacobs and Huang, 2019). When Notch signalling was inhibited from 18 hpf to 30 hpf, we saw two distinct patterning phenotypes depending on the anterior-posterior positioning in the spinal cord (Fig. 2A). In the anterior region, we saw a vast increase in the number of V3 cells with little effect on KA” cells. By contrast, the posterior region showed a dramatic increase in the number of KA” cells at the expense of V3 cells. Zebrafish spinal cord development occurs along the length of the anterior-posterior axis, so the more anterior tissue is “older” than the posterior tissue. At the point of Notch inhibition, the anterior tissue had already completed the wave of KA” differentiation, while the more posterior tissue had not. While V3 cells could differentiate from LFP progenitors in the anterior tissue, the “younger” posterior tissue differentiated into KA” cells and depleted the pool of progenitors to prevent V3 differentiation. Interestingly, in the intermediate region, both cell types can be seen, including some cells expressing both *tal2* and *sim1*, suggesting a mixed fate (Fig. 2A). In addition, we combined the *vglut2a:EGFP* reporter, which labels the same V3 cell population as the *vglut2a:dsRed* reporter (Fig. S2), with a *nkx2.2a:NLS-mCherry* transgenic line (Zhu et al., 2019) to visualise the V3 cells alongside LFP progenitors. As mCherry is a stable protein, it perdures within the cell after the promotor has been switched off. We were able to visualise over 90% of *vglut2a:EGFP^+^* cells that still expressed mCherry at 48 hpf (Fig. 2B, C), a time point when only 17% of the *sim1^+^* V3 cells still expressed *nkx2.2a* transcript (Fig. S1D). Taken together, our data support the model that KA” and V3 cells come from a common LFP lineage in a sequential manner.

**Figure 2.**
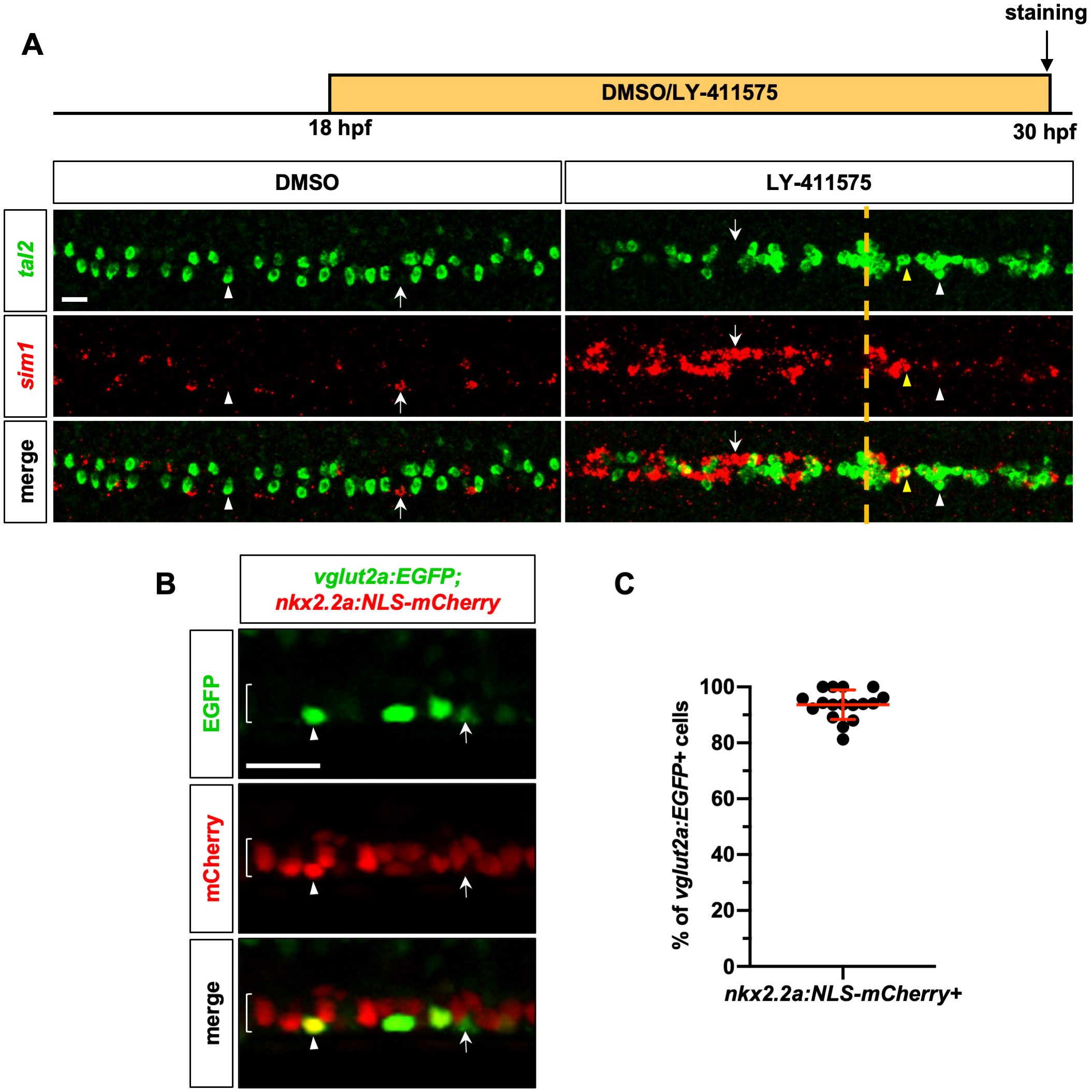
V3 interneurons are derived from the *nkx2.2a* expressing LFP lineage. (A) Wild type embryos were incubated in either DMSO or LY-411575 from 18 hpf until fixation at 30 hpf. Whole-mount double fluorescent in situ hybridisation was performed for *tal2* and *sim1* and imaging was from the dorsal view. The experimental procedure is shown on the top. White arrows and arrowheads highlight *sim1^+^* V3 cells and *tal2^+^* KA” cells, respectively. Yellow arrowheads denote a *sim1^+^tal2^+^* cell. Dotted yellow line denotes the estimated point of development the drug took effect. *n* = 6 embryos per condition. (B) *vglut2a:EGFP; nkx2.2a:NLS-mCherry* embryos were imaged at 48 hpf. Arrowheads highlight an EGFP^+^mCherry^+^ cell. Arrows highlight a *vglut2a:EGFP* positive cell that is negative for *nkx2.2a:NLS-mCherry*. Bracket denotes the LFP domain. *n* = 17 embryos. (C) The number of *vglut2a:EGFP; nkx2.2a:NLS-mCherry* double positive cells displayed as a percentage of the total *vglut2a:EGFP* positive cell population. Each data point represents the percentage of double positive cells in one embryo. The image shown in (B) is an example of one of these embryos. Data are plotted with mean ± SD. Scale bars: 20 μm.

### The cell types of the LFP can be distinguished by their Notch signalling profiles

As we have shown that the attenuation of Notch signalling causes spontaneous differentiation into either KA” or V3 cells depending on the time of inhibition (Fig. 2A) (Huang et al., 2012), we next utilised our previously described Notch signalling reporter (Jacobs and Huang, 2019) to analyse the levels of Notch response in the cell populations of the LFP domain. When visualised through immunostaining for GABA in the context of the *her12:Kaede* reporter, we discovered GABA^+^ KA” cells had a distinct lack of total Notch response by 24 hpf. They could be clearly identified by the lack of *her12:Kaede* expression, and their characteristic medial location in the LFP domain (Fig. 3A). Interestingly, there were cells with a similarly low level of Notch response but not labelled by GABA (Fig. 3A). We hypothesised that these GABA^-^Kaede^low^ cells were likely the earliest born V3 interneurons. To test this possibility, we crossed the *vglut2a:dsRed* transgenic reporter with *her12:Kaede* to visualise Notch response. This allowed us to identify dsRed^+^ V3 interneurons alongside KA” cells with the characteristic low level of *her12:Kaede* expression, and the remaining LFP progenitor cells (Fig. 3B). As the Kaede protein perdures over the timepoints we analysed, we can use the level of Kaede fluorescence to estimate the total amount of Notch signalling each cell received, as *her12* expression is a readout for Notch activity. Kaede fluorescence thus represents the combination of early signalling response and ongoing expression. Through quantification of the total levels of Kaede fluorescence at 32 hpf, we discovered the three cell populations had discrete Notch signalling profiles (Fig. 3C). KA” cells displayed low, but not absent, levels of Notch response. V3 cells showed higher levels of total Notch response compared to KA” cells, but still significantly less than the high levels present in LFP progenitor cells. This result suggests that the early, homogenous population of LFP progenitors receive a low level of Notch signalling (Notch^LO^) before a subset switches off their response to differentiate into Notch^OFF^ KA” interneurons (Fig. 3D). Later another subset switches off their response to differentiate into Notch^OFF^ V3 cells, while LFP progenitors maintain high levels of Notch signalling (Notch^HI^) (Fig. 3D). Therefore, KA” cells, V3 cells, and LFP progenitor cells receive increasingly higher amounts of total Notch response.

**Figure 3.**
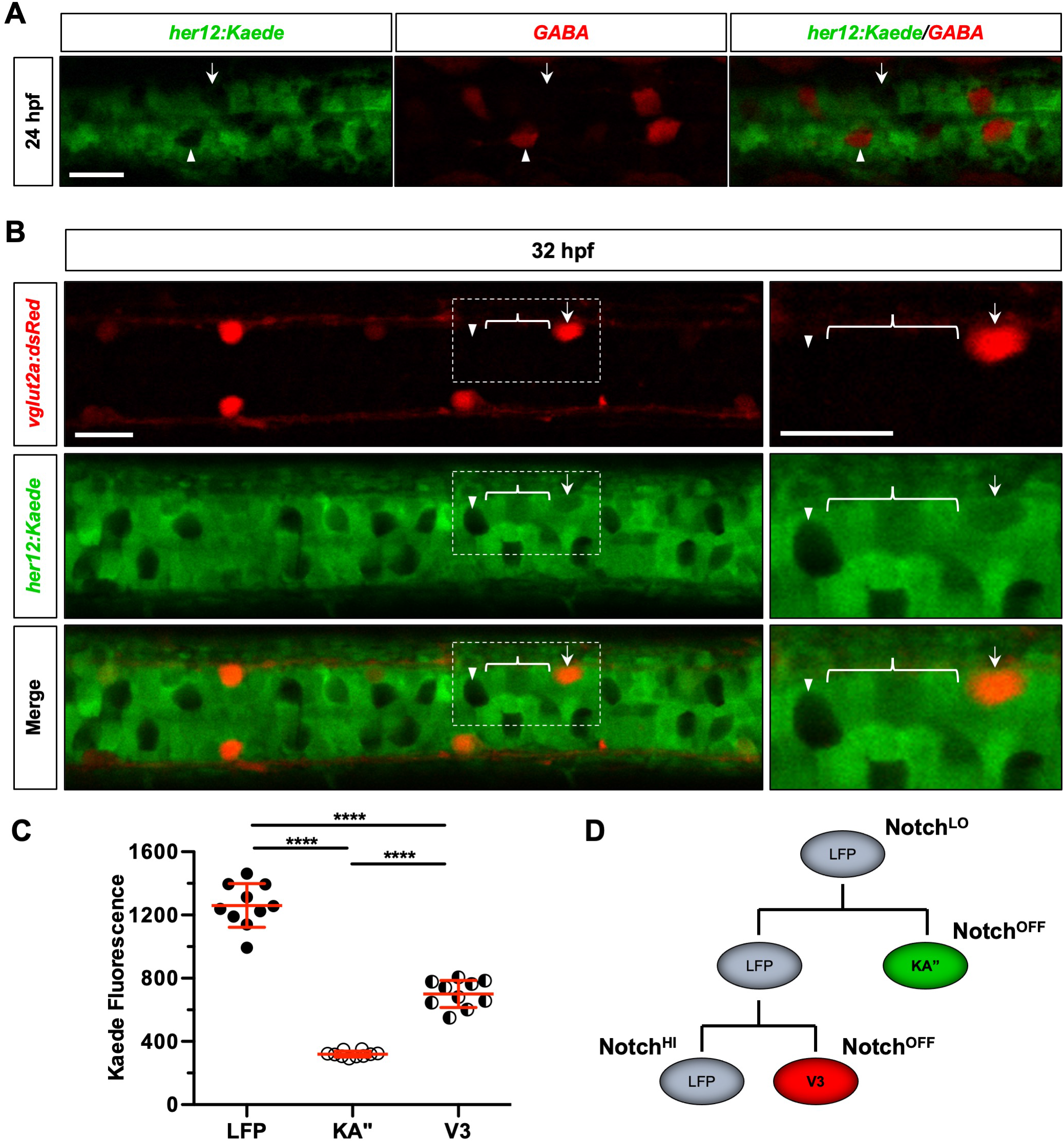
Different cell types in the LFP domain display discrete Notch signalling profiles. (A) *her12:Kaede* embryos were fixed at 24 hpf and immunohistochemistry for GABA was performed to label KA” cells (arrowheads). Embryos were imaged from the dorsal view. Arrows highlight potential V3 interneurons due to the lack of GABA and *her12:Kaede* expression. *n* = 4 embryos. (B) *her12:Kaede; vglut2a:dsRed* embryos were live imaged at 32 hpf from the dorsal view. Arrowheads highlight a KA” cell, based on the data shown in (A). Arrows highlight a *vglut2a:dsRed^+^* V3 cell. Brackets denote area defined as LFP progenitors. Expanded views of the boxed regions are shown on the right. (C) Kaede fluorescence from the experiment shown in (B) was measured for 10 examples of KA”, V3 or LFP cells per embryo. Each data point represents the average Kaede fluorescence of one embryo. Data are plotted with mean ± SD. Statistics: Kruskal-Wallis, medians vary significantly (****), Mann-Whitney *U* test (shown). Asterisks representation: p-value<0.0001 (****). *n* = 10 embryos. (D) Schematic showing the predicted Notch signalling dynamics during cell fate specification in the LFP. Scale bars: 20 μm.

These distinct total Notch signalling profiles do not describe whether the LFP progenitors receive sustained Notch signalling over time, or if they are exposed to different doses dependent on what fate they will become. The total Notch signalling profiles reflect the cumulative Notch response that occurs in the cell and its progenitors up to 32 hpf. To determine Notch response in a defined time window, we leveraged the photoconvertibility of the *her12:Kaede* reporter. Briefly, while mature Kaede protein can be photoconverted from green to red fluorescence, the newly transcribed and translated Kaede after the photoconversion will be in the green state. This allows us to visualise Notch response that occurs during a specific time window (Jacobs and Huang, 2019). Combining *her12:Kaede* with the *vglut2a:dsRed* reporter, we analysed the Notch signalling profiles of each cell population in the LFP during 24 hpf and 32 hpf by photoconverting embryos at 24 hpf (Fig. 4A, B). During this time period, KA” cells received no Notch signalling (Fig. 4C), which is to be expected as KA” determination and differentiation is complete by 24 hpf (Fig. 1E). On the other hand, V3 interneurons continued to receive Notch signals during this time (Fig. 4C), demonstrating an extended duration of Notch response in comparison to KA” cells. Finally, LFP progenitor cells received a higher level of Notch signalling during this time period (Fig. 4C). Together, these results show that not only do the distinct cell populations of the LFP domain receive different total levels of Notch signalling, but this total is achieved through differing durations of Notch response (Fig. 4D).

**Figure 4.**
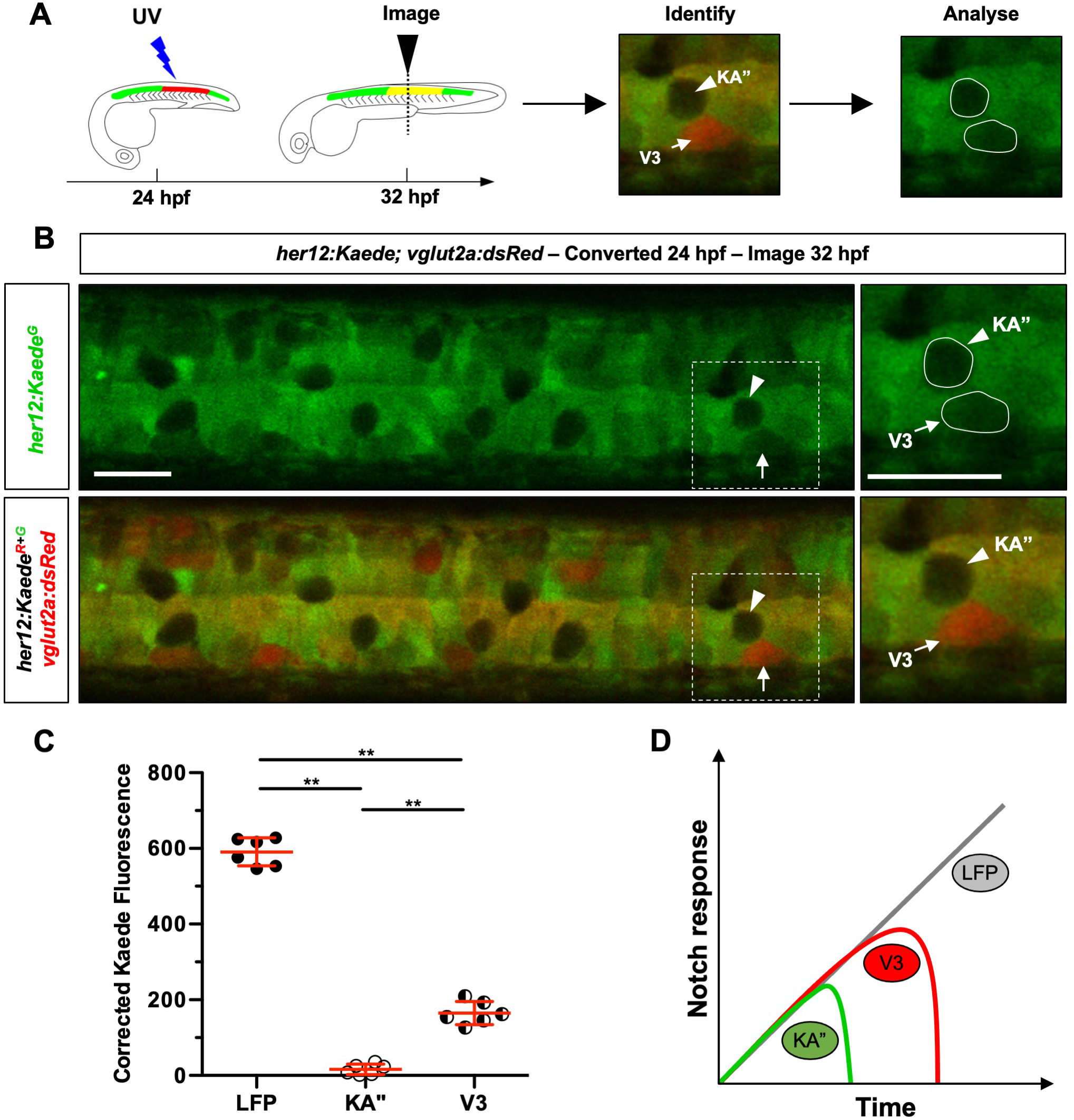
Different cell types in the LFP domain show different durations of Notch signalling. (A) Schematic representation of the experimental design for (B). (B) A section of the spinal cord of *her12:Kaede; vglut2a:dsRed* embryos was photoconverted by UV light at 24 hpf and the Kaede^green^ fluorescent profiles of the different cell types were analysed from the dorsal view at 32 hpf. Cell types were identified in the same manner as Fig. 3. Expanded views of the boxed regions are shown on the right. (C) Kaede fluorescence from the experiment shown in (B) was measured for 10 examples of KA”, V3 or LFP cells per embryo. The average level of unconverted Kaede^green^ fluorescence was calculated at 24 hpf, directly after photoconversion and subtracted from the value recorded at 32 hpf to generate the corrected Kaede fluorescence values displayed. Each data point represents the average corrected Kaede fluorescence of one embryo. Data are plotted with mean ± SD. Statistics: Kruskal-Wallis, medians vary significantly (****), Mann-Whitney *U* test (shown). Asterisks representation: p-value<0.01 (**). *n* = 6 embryos. (D) Schematic representation of the different durations of Notch signalling each cell type in the LFP domain receives. Scale bars: 20 μm.

### The duration of Notch signalling is instructive in LFP fate determination

To interrogate the importance of duration of signal versus total level, we artificially sustained Notch signalling at a high level in the ventral spinal cord using a *ptc2:Gal4; UAS:NICD* (referred to as *ptc2:NICD*) transgenic line. Since we have previously shown that Hh signalling is positively regulated by Notch signalling (Jacobs and Huang, 2019) and *ptc2* is a direct downstream target of Hh response, expressing NICD under the control of the *ptc2* promoter (*ptc2:NICD*) would form a positive feedback loop to sustain Notch signalling. Indeed, *ptc2:NICD* embryos showed elevated level of *her12* expression, indicative of enhanced Notch activation (Fig. 5A). Consequently, sustained Notch signalling prevented V3 and KA” differentiation in the LFP, leading to an expansion of *nkx2.2a^+^* LFP progenitors (Fig. 5B, C). We then combined this sustained Notch^ON^ system with the *hsp:dnMAML-GFP* (referred to as *hsp:dnMAML*) transgenic line, a heat inducible terminator of Notch signalling (Zhao et al., 2014). This allowed us to selectively switch off the sustained, high level Notch signalling through heat shock at specific time points. Generating *ptc2:NICD; hsp:dnMAML* embryos resulted in siblings of wild-type, *hsp:dnMAML, ptc2:NICD* and *ptc2:NICD; hsp:dnMAML* genotypes, allowing us to examine all conditions as internal controls.

**Figure 5.**
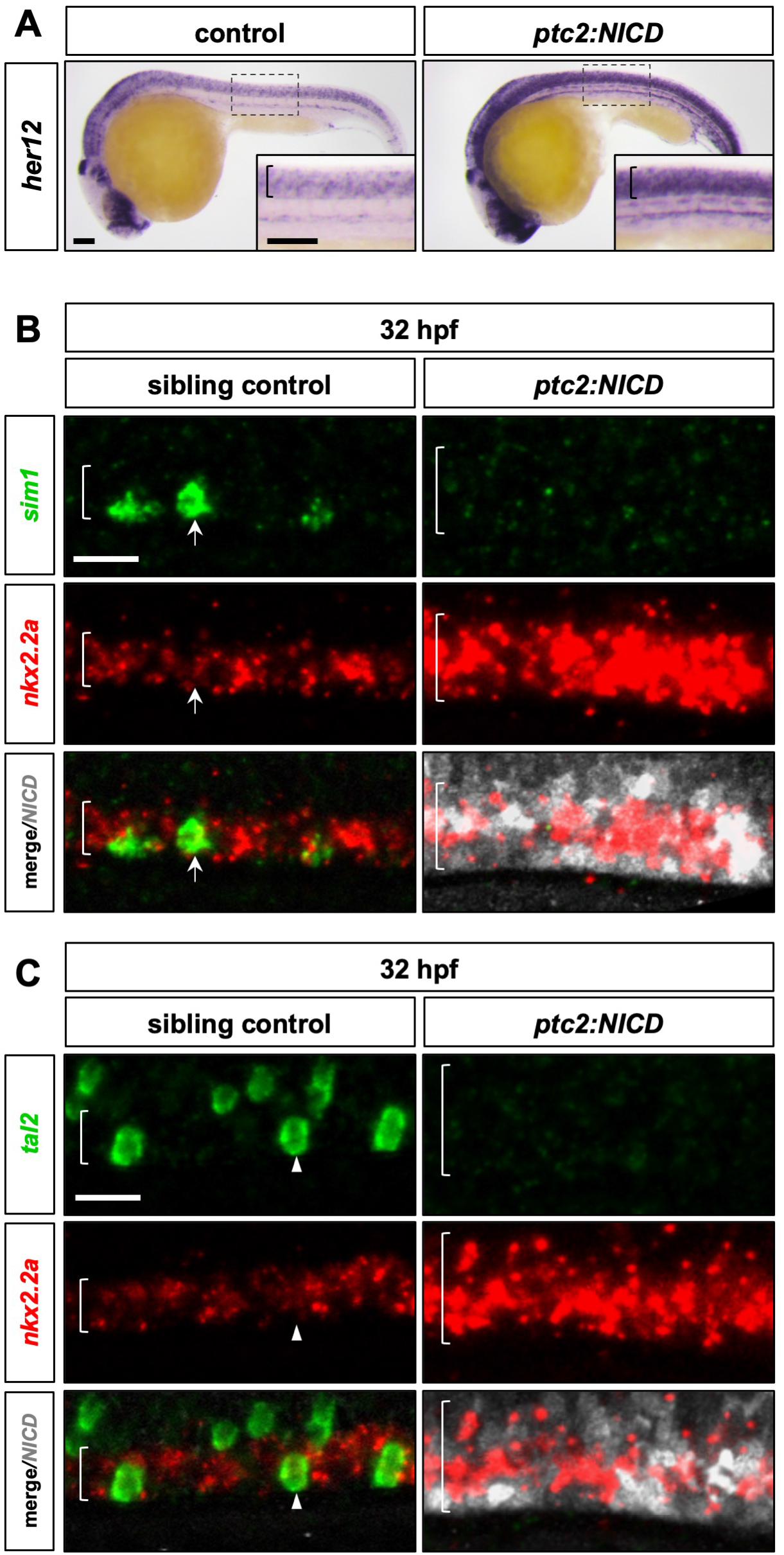
Sustained Notch signalling prevents KA” and V3 differentiation. (A) Whole-mount in situ hybridisation for *her12* was performed on *ptc2:Gal4; UAS:NICD* (shown as *ptc2:NICD*) and their wild type sibling embryos at 24 hpf. Fixed embryos were imaged from the lateral view. *n* = 15 embryos per condition. Bracket shows the spinal cord. (B, C) *ptc2:Gal4; UAS:NICD* (shown as *ptc2:NICD*) and their wild type sibling embryos were fixed at 32 hpf and whole-mount double fluorescent in situ hybridisation for either *sim1* (B) or *tal2* (C) and *nkx2.2a* alongside immunohistochemistry for Myc to label Myc tagged NICD expression. Fixed embryos were imaged from the lateral view. Brackets denote the dorsoventral extent of the *nkx2.2a^+^* domain. Arrows in (B) highlight a *sim1^+^* V3 cell. Arrowheads in (C) highlight a *tal2^+^* KA” cell. *n* = 4 embryos per condition. Scale bars: (A) 100 μm; (B, C) 20 μm.

These embryos were heat shocked at either 14 hpf or 24 hpf, and then KA” and V3 cell fates were analysed at 36 hpf (Fig. 6A). As expected, inhibiting Notch signalling with *hsp:dnMAML* at 14 hpf resulted in a dramatic increase in the number of KA” cells at 36 hpf, at the expense of V3 cells (Fig. 6B, D). Interestingly, KA” cell numbers in *ptc2:NICD; hsp:dnMAML* embryos were similar to the wild-type siblings while there was an almost complete absence of V3 cells (Fig. 6B, D). Despite the high levels of Notch signalling generated through the *ptc2:NICD* line (Fig. 5A), switching off Notch signalling at 14 hpf causes KA” differentiation and does not allow for V3 fate.

**Figure 6.**
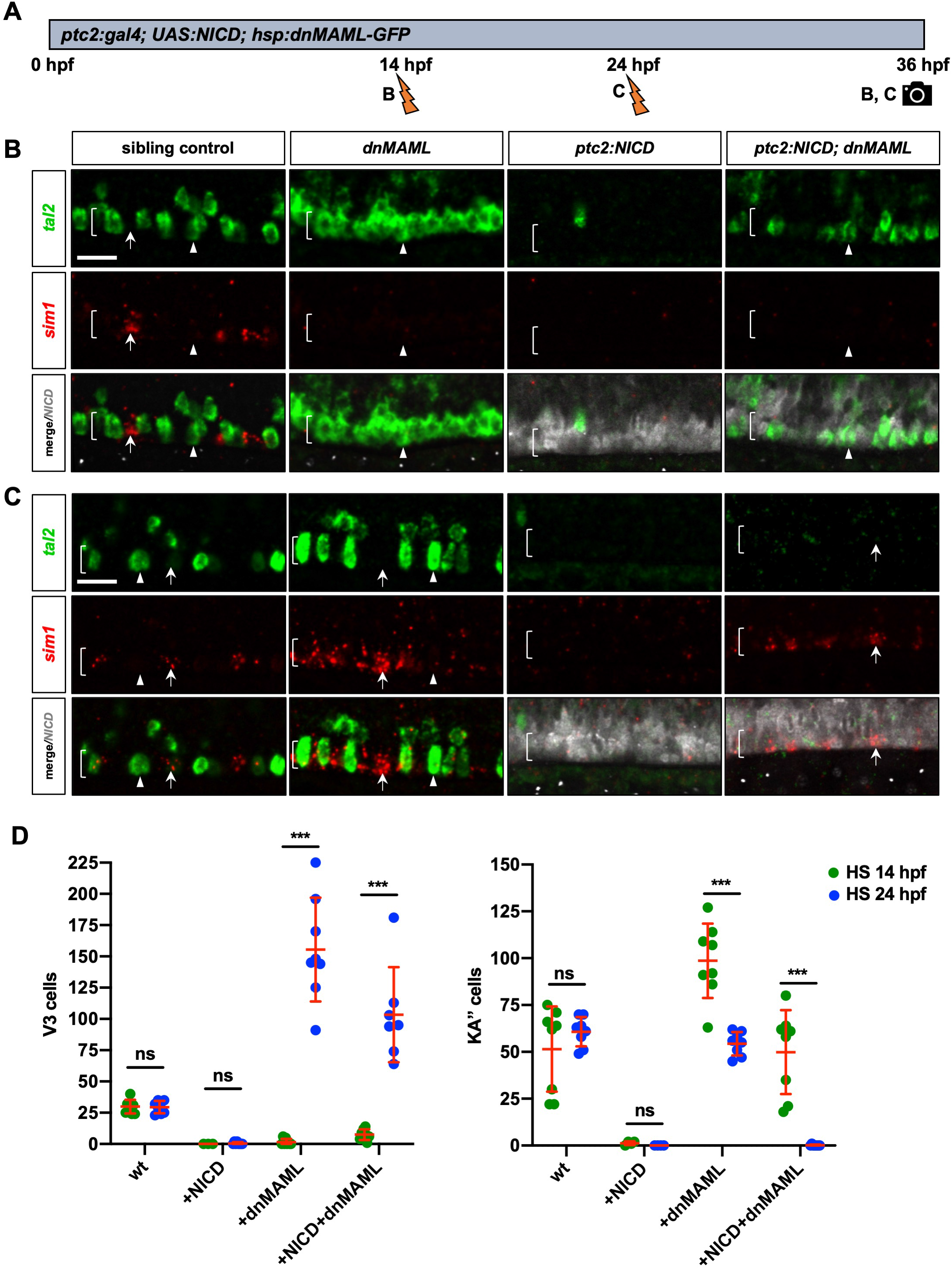
The duration of Notch signalling is instructive in LFP cell fate determination. (A) Schematic representation of the experimental design for (B) and (C). (B, C) *hsp:dnMAML-GFP* (shown as *dnMAML*), *ptc2:Gal4; UAS:NICD* (shown as *ptc2:NICD*), *ptc2:NICD; hsp:dnMAML-GFP* and their wild type sibling embryos were heat shocked at either 14 hpf (B) or 24 hpf (C) then fixed at 36 hpf. Whole-mount double fluorescent in situ hybridisation for *tal2* and *sim1* was performed alongside immunohistochemistry for Myc to label Myc tagged NICD expression. Fixed embryos were imaged from the lateral view. Arrowheads highlight *tal2^+^* KA” cells. Arrows highlight *sim1^+^* V3 cells. Brackets denote the extent of the LFP domain. *n* = 4 embryos per condition. (D) Embryos from similar experiments as in (B) and (C) were stained with *sim1* or *tal2* at 36 hpf. The number of V3 (*sim1^+^*) or KA” (*tal2^+^*) cells was quantified under each condition. Each data point represents the number of cells in one embryo. Data are plotted with mean ± SD. Statistics: Mann-Whitney *U* test. Asterisks representation: p-value<0.001 (***); ns: not significant. *n* = 8 embryos per condition. Scale bars: 20 μm.

Inhibition of Notch signalling at 24 hpf caused the opposite effect. In *hsp:dnMAML* embryos heat shocked at 24 hpf, there was a dramatic increase in the number of V3 cells at 36 hpf, while the KA” population was unaffected (Fig. 6C, D). This is consistent with our earlier results (Fig. 2A), as KA” differentiation has been completed prior to 24 hpf. In *ptc2:NICD; hsp:dnMAML* embryos heat shocked at 24 hpf, we saw a similar increase in the number of V3 cells but, critically, no KA” cells were present (Fig. 6C, D). This demonstrates that sustaining Notch signalling until 24 hpf prevents KA” differentiation and leads exclusively to V3 cell fate. Taken together, our results suggest that the duration of Notch signalling is instructive during fate determination in the LFP.

We next switched off the sustained Notch signalling at 18 hpf (Fig. S3A), a time point near the end of KA” differentiation and right at the beginning of V3 differentiation (Fig. 1D-E). In *hsp:dnMAML* embryos, there were a similar number of differentiated cells compared to the wild-type siblings, with a slight bias towards KA” cells over V3 cells (Fig. S3B). Intriguingly, in *ptc2:NICD; hsp:dnMAML* embryos, we visualised both KA” and V3 cells, but with a bias toward V3 (Fig. S3B). To examine whether the stalled progenitors created by the sustained Notch signalling still held their potential to differentiate at later stages, we heat shocked the embryos at 36 hpf and analysed them at 48 hpf (Fig. S3A). Impressively, the *ptc2:NICD; hsp:dnMAML* embryos were still able to generate a similar number of V3 cells compared to both the *hsp:dnMAML* and wild-type siblings (Fig. S3C). As expected, no KA” cells were present in *ptc2:NICD; hsp:dnMAML* after heat shock at 36 hpf as Notch signalling had been sustained throughout KA” differentiation (Fig. S3C). These Notch^ON^-Notch^OFF^ experiments demonstrate that the duration of Notch signalling, irrespective of the level, can be used to describe the different waves of differentiation in the LFP – progenitors that differentiate into KA” cells are exposed to a short duration of Notch signalling that ends after 18 hpf but before 24 hpf, while progenitors that differentiate into V3 cells are exposed to a longer duration of Notch signalling that can continue past 36 hpf.

### Jag2b regulates KA” and V3 formation within their normal differentiation timetable

As the LFP domain begins as a seemingly homogenous population of progenitors and the differentiated neurons appear in a stochastic “salt and pepper” pattern, what could be causing direct neighbours to have distinct Notch response profiles? As the Notch ligand *jag2b* has been previously implicated in KA” cell differentiation (Yeo and Chitnis, 2007), we postulated that Jag2b functions to maintain LFP progenitors to limit the differentiation of KA” and V3 cells. Expression analysis at 24 hpf revealed that *jag2b* was expressed strongly in the domain dorsal to *tal2^+^* KA” cells (Fig. 7A), likely corresponding to the motor neuron progenitor domain as previously reported (Yeo and Chitnis, 2007). Similarly, *jag2b* was also not expressed in *sim1^+^* V3 cells at 30 hpf (Fig. 7B). To analyse whether Jag2b is mediating Notch signalling in the LFP, we used morpholino knockdown to determine how the loss of *jag2b* affects LFP patterning. We injected a specific morpholino targeting *jag2b* (*jag2b^MO^*) (Liu et al., 2007) to knockdown *jag2b* and then quantified the number of KA” and V3 cells at 20 hpf and 30 hpf (Fig. 7C-E). Since *tal2* labels both KA” and the slightly dorsal KA’ cells, we used the KA” specific marker *urp1* (Quan et al., 2015) (Fig. S4) instead to ensure reliable counting of KA” cells. Knockdown of *jag2b* lead to a 20% increase in the number of KA” cells at 30 hpf (Fig. 7C, D). Interestingly, at 20 hpf this increase was not present (Fig. 7D). Similarly, *jag2b* knockdown also caused a striking 52% increase in the number of V3 cells present at 30 hpf, while the number of cells at 20 hpf was unaffected (Fig. 7C, E). This result suggests that Jag2b-mediated Notch signalling in LFP progenitors acts to prevent their differentiation into KA” and V3 cells. Since the loss of Jag2b did not result in precocious KA” and V3 formation before their normal differentiation timetable, there is likely a Jag2b-independent mechanism controlling the timing of KA” and V3 differentiation.

**Figure 7.**
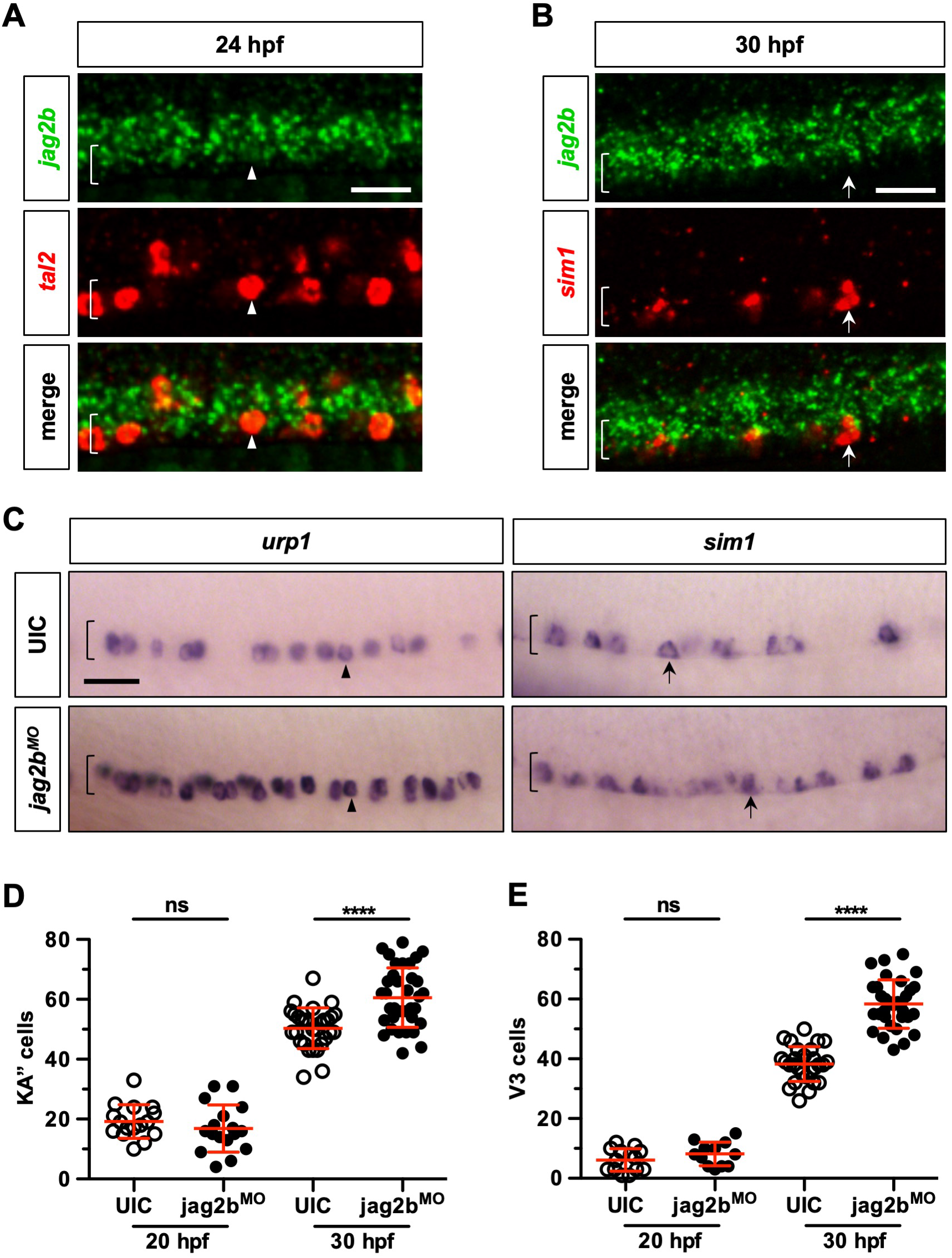
Knockdown of *jag2b* promotes V3 and KA” differentiation. (A, B) Whole-mount double fluorescent in situ hybridisation was performed on wild type embryos for *jag2b* together with either *tal2* at 24 hpf (A), or *sim1* at 30 hpf (B). Brackets denote the extent of the LFP domain. (C) Whole-mount in situ hybridisation for either *urp1*, to label KA” cells (arrowheads), or *sim1*, to label V3 cells (arrows), was performed in *jag2b*^MO^ injected fish and uninjected controls (UIC). Fixed embryos were imaged from the lateral view. Brackets denote the extent of the LFP domain. (D, E) The number of KA” or V3 cells were counted from the experiment shown in (C) as well as at 20 hpf. Each data point represents the number of cells in one embryo. Data are plotted with mean ± SD. Statistics: Mann-Whitney *U* test. Asterisks representation: p-value<0.0001 (****); ns: not significant. *n* = 14-36 embryos per staining. Scale bars: 20 μm.

## DISCUSSION

The mechanisms by which neural progenitor cells switch from progenitor states to differentiated neurons have long been an area of much interest. Here, using the zebrafish lateral floor plate as a model, we demonstrate that the duration of Notch signalling plays an instructive role in temporal cell fate specification. First, common LFP progenitors generate KA” and V3 interneurons in a sequential manner. Second, different cell types in the LFP domain display discrete Notch signalling profiles. Third, using a conditional Notch^ON^-Notch^OFF^ system, we demonstrate a duration-specific role for Notch signalling in cell fate decisions during spinal cord patterning.

### Differentiation dynamics and lineage relationships in the ventral spinal cord

Neural differentiation in the brain has previously been described as waves, with different cell fates appearing during specific time windows (Agirman et al., 2017; Noctor et al., 2004). Here, we describe a similar temporal process of cell fate specification in the most ventral domain of the zebrafish spinal cord. After the initial, early, wave of KA” differentiation, the progenitors of the LFP then enter a prolonged period of V3 differentiation. This long wave of V3 differentiation is characterised by cells first appearing at the lateral edge of the LFP, before slowly occupying the more medial locations. Intriguingly, these waves of differentiation finish without filling the entire LFP with either KA” or V3 interneurons, suggesting a late maintained pool of progenitors. These remaining *nkx2.2a^+^* LFP progenitors likely contribute to perineurial glia to regulate and support motor axons (Kucenas et al., 2008). Unlike KA” cells, which do not co-express the progenitor marker *nkx2.2a* at any timepoint analysed (Huang et al., 2012), V3 cells go through a stage where they express both the fate marker *sim1* and the progenitor marker *nkx2.2a*. Could this be a sign of higher levels of plasticity in the early stages of fate determination? More likely, the well-studied markers for KA” fate, *tal2* and *gata2* specifically (Yang et al., 2010; Yang et al., 2020), are comparably later markers than *sim1* in V3 fate. Also, as only a minority of *sim1* expressing V3 cells express the terminal marker HuC, it is possible that V3 fate determination is a longer process when compared to the early-born KA” fate.

Previously, we described a “salt-and-pepper” organisation of the LFP domain which tends to leave 1-4 cell gaps of progenitors between the differentiated KA” interneurons (Huang et al., 2012). Here, we show that a subset of the progenitors that do not become KA” interneurons go on to differentiate into V3 cells. Similar to KA” differentiation, the pattern in which they arise is stochastic. Inhibition of Notch signalling at a mid-stage between the peaks of KA” and V3 differentiation forces most LFP cells to become either population, depending on their position along the anterior-posterior axis. This demonstrates that all LFP progenitors have the potential to differentiate into either KA” or V3 but receive an extrinsic timing cue in order to control which fate they acquire, or whether they remain a progenitor. This cue could be Hh signalling, as KA” cells lose their Hh response while the neighbouring LFP progenitors do not (Huang et al., 2012). However, as we have demonstrated that the Hh response in the spinal cord is controlled through Notch signalling (Jacobs and Huang, 2019), and due to the canonical role of Notch signalling in progenitor maintenance (Louvi and Artavanis-Tsakonas, 2006), it is likely that Notch signalling is providing the ultimate signal controlling fate determination in the LFP, though we cannot rule out other external signals.

### Discrete Notch signalling profiles during spinal cord development

As KA” cells show differential Hh response compared to their LFP neighbours (Huang et al., 2012), we expected a similar pattern of Notch response. Our Notch signalling reporter *her12:Kaede* allows us to visualise the differential Notch response of both KA” and V3 cells and reveals a different pattern compared to Hh response. KA” cells attenuate their Hh response when they differentiate but have high levels of Hh signalling history, showing that the progenitors do previously have high levels of Hh response (Huang et al., 2012). This is not the case for Notch signalling. KA” cells have very low levels of total Notch response, LFP progenitors have a much higher total level, while V3 interneurons are somewhere in between. This suggests that the early, homogeneous population of LFP progenitors are maintained by a low level of Notch signalling, prior to requiring a higher level to prevent aberrant differentiation during KA” fate determination.

Through quantifying our PHRESH analysis, we reveal that progenitors that differentiate into KA” cells receive the shortest duration of Notch signalling before attenuation, while progenitors that differentiate into V3 cells receive a longer duration before also losing their response when they differentiate. By contrast, LFP progenitors maintain active Notch signalling throughout the timepoints analysed, further highlighting the requirement for an upregulation of Notch signalling in progenitors during the different waves of differentiation. These different cell fates being exposed to different durations of Notch signal is reminiscent of the “temporal adaptation” model of Hh signalling described in ventral spinal cord patterning, which suggests the most ventral fates require both a higher level and a longer duration of Hh signalling (Dessaud et al., 2007). Here, it is likely the different total levels of Notch response observed are simply a function of the duration, but further studies are required to determine the linearity of Notch signalling in this system. The level and duration of Notch signalling may also be a general controller of spinal cord neurogenesis, since we have observed differential Notch response of other differentiated neurons, such as KA’ cells, compared to their direct neighbours, as suggested by a previous study (Shin et al., 2007).

### The duration of Notch signalling is an instructive cue in fate determination

The loss of Notch response causes spontaneous differentiation in the spinal cord, as shown by our loss-of-function experiments (Fig. 2A, 6B) alongside other studies (Appel et al., 2001; Huang et al., 2012; Park and Appel, 2003). Which fate the progenitors acquire is dependent on the time Notch signalling is inhibited (Huang et al., 2012; Jacobs and Huang, 2019; Yeo and Chitnis, 2007). Using the Gal4-UAS system, which is known to drive high levels of ectopic expression (Scheer and Campos-Ortega, 1999), we develop a Notch^ON^-Notch^OFF^ system to sustain Notch signalling at a high level in the ventral spinal cord, then systematically attenuate the Notch response throughout spinal cord patterning to interrogate the importance of Notch signalling duration versus level.

First, a sustained, high level of Notch signalling stalls progenitor cells and prevents differentiation into either early-born KA” or later-born V3 cells. This demonstrates that attenuation of Notch signalling is required for neuronal differentiation. Second, despite high levels of signalling, early termination of Notch signalling can only generate KA” fate. In order to acquire V3 fate, LFP progenitors must receive Notch signals until at least 18 hpf. Interestingly, at this time point progenitors can acquire either fate, irrespective of their anterior-posterior position, suggesting a potential cell-fate specific mechanism downstream of, or in parallel to, Notch signalling. This establishes an instructive role for an extended duration of Notch signalling, rather than simply a higher threshold of total level, in the acquisition of V3 over KA” cell fate. It appears to be the timing at which Notch signalling is attenuated in LFP progenitor cells that provides the instructive cue, as cells rapidly differentiate into either KA” or V3 cells dependent upon the timing of Notch inhibition. Finally, the duration of Notch signalling required for V3 cell identity is more generous than for KA” differentiation. Stalled progenitors retain their potential to become V3 cells for an extended period, while KA” cells can only be generated within a tight, early time window. This is likely due to lineage constraints – V3 differentiation occurs directly after the tight window of KA” differentiation, and then has a much longer time window, allowing for any late differentiation event to generate V3 cells. This could be part of the mechanism that causes the long-wave of V3 differentiation when compared to KA” differentiation.

### Two parallel roles for Notch signalling in the LFP

What is the mechanism by which the initial, homogeneous population of LFP progenitor cells selectively attenuate their Notch response? It is unlikely that the early progenitor cells have an intrinsic program to become either KA” or V3 cells, as late inhibition of Notch signalling causes V3 differentiation with no KA” differentiation. Indeed, based on the upregulation of Notch signalling in cells that do not become KA” cells, it is likely that the default fate of the early progenitor cells is KA” fate and exposure to Notch signalling is what extends progenitor state to allow for V3 differentiation at a later stage. We observe a time point at which, when Notch signalling is switched off, LFP progenitors can differentiate into either KA” or V3 cells. How do they decide? Also, establishing the correct number of neurons is integral for CNS function. Does the same mechanism that controls the cell fate decision also mediate the expansion of each cell population?

General inhibition of Notch signalling from an early stage leads to depletion of the progenitor pool through spontaneous differentiation into the earliest-born motor neurons (Appel et al., 2001). Intriguingly, consistent with previous reports (Yeo and Chitnis, 2007), *jag2b* knockdown results in the opposite phenotype – an increase in the number of both KA” and V3 cells. This result suggests that Notch signalling mediated through Jag2b plays a general role, not a cell fate specific one, in preventing neural differentiation of LFP progenitors. Importantly, it reveals that there is likely a mechanism in place that prevents runaway differentiation upon the loss of *jag2b*, maintaining the specific timetable of Notch signalling that results in KA” or V3 fate. As the increased differentiation of both KA” and V3 cells upon the loss of *jag2b* occurs during their regular time windows, this parallel mechanism must be able to maintain Notch response independent of Jag2b. It has been shown that domain-specific expression of Notch ligands leads to Notch response that maintains different progenitor domains (Marklund et al., 2010), which suggests that different ligand-mediated Notch signals could be involved in a fate-specifying role. It is plausible to suggest that two parallel mechanisms of Notch signalling are controlling patterning in the LFP – one that decides between KA” and V3 fate and one, mediated by Jag2b, that maintains LFP progenitors.

In summary, we describe the unique differentiation dynamics of the lateral floor plate lineage during spinal cord patterning in zebrafish. We show that the duration of Notch signalling functions as an instructive cue in temporal cell fate determination, with extended durations leading to exclusively late-born fates. Future work on different Notch ligands will likely provide new insights on how the duration of Notch signalling is precisely controlled to achieve reproducible pattern formation.

## MATERIALS AND METHODS

### Zebrafish strains

All zebrafish strains used in this study were maintained and raised under standard conditions. All procedures were conducted in accordance with the principles outlined in the current Guidelines of the Canadian Council on Animal Care. All protocols were approved by the Animal Care Committee at the University of Calgary (#AC17-0128 and #AC21-0102). The transgenic strains used were: *Tg(gata2:GFP)la3* (Traver et al., 2003), *TgBAC(her12:Kaede)ca106* (Jacobs and Huang, 2019), *Tg(hsp:dnMAML-GFP)ca113, Tg(UAS:NICD)kca3* (Scheer and Campos-Ortega, 1999), *Tg(nkx2.2a:NLS-mCherry)ca114, Tg(UAS:NTR-mCherry)c264* (Davison et al., 2007), *TgBAC(ptc2:Gal4)ca112, Tg(slc17a6b:dsRed)nns9 (vglut2a:dsRed)* (Miyasaka et al., 2009), and *Tg(slc17a6b:EGFP)zf139 (vglut2a:EGFP)* (Bae et al., 2009).

### Generation of transgenic lines

The *hsp:dnMAML-GFP* (Zhao et al., 2014) and *nkx2.2a:NLS-mCherry* (Zhu et al., 2019) transgenic lines were generated by standard Tol2-mediated transgenesis. To generate the *ptc2:Gal4* transgenic line, BAC clone zC226H23 from the CHORI-211 library that contains the *ptc2* genomic region was selected for bacteria-mediated homologous recombination following standard protocols (Bussmann and Schulte-Merker, 2011). zC226H23 contains 150 kb upstream and 20 kb downstream of flanking sequence with potential regulatory elements. First, a cassette containing two opposite facing Tol2 arms and an ampicillin resistance gene was recombined into the BAC vector of zC226H23. Then a cassette containing the transcriptional activator Gal4-VP16 (Sadowski et al., 1988) with the kanamycin resistance gene was recombined into the BAC to replace the first exon of the *ptc2* gene. Successfully modified BAC constructs were confirmed by PCR analysis. The recombinant *ptc2:Gal4* construct was then co-injected with *tol2* transposase mRNA into *UAS:NTR-mCherry* embryos at the one-cell stage. Stable transgenic lines were established by screening for mCherry expression in F1 embryos from injected founders.

### Morpholino injection

Morpholino oligonucleotides (Gene Tools, LLC) targeting *jag2b* (5’-TCCTGATACAATTCCACATGCCGCC-3’) (Liu et al., 2007) were injected at 0.4-0.6 mM into one-cell stage embryos with 1 nl per embryo. Injected embryos were fixed at appropriate stages for in situ analysis.

### In situ hybridisation and immunohistochemistry

All whole-mount in situ hybridisation and immunohistochemistry in this study were performed using standard protocols. We used the following antisense RNA probes: *her12*, *jag2b*, *nkx2.2a*, *sim1*, *tal2*, and *urp1*. For double fluorescent in situ hybridisation, both dinitrophenyl (DNP) and digoxigenin (DIG) labelled probes were used with homemade FITC and Cy3 tyramide solutions (Vize et al., 2009). For immunohistochemistry, we used the following primary antibodies: mouse polyclonal antibodies to HuC (1:1000, Invitrogen) and Myc (1:1000, DSHB), and rabbit polyclonal antibodies to GABA (1:500, Sigma) and RFP (1:1000, MBL). The appropriate Alexa Fluor-conjugated secondary antibodies were used (1:500, Thermo Fisher) for fluorescent detection of antibody staining and Draq5 (1:10,000, Biostatus) was used for nuclear staining. To visualise *vglut2a:dsRed* alongside *sim1*, whole mount immunohistochemistry was carried out first using standard procedure that included RNase inhibitor (1:100, Roche) during the blocking steps and used a primary antibody for RFP, followed by standard fluorescent in situ hybridisation using a *sim1* DIG labelled probe.

### Heat shock and drug treatment

To induce expression from the heat shock promoter, embryos at the relevant stage were placed in a 2 ml micro-centrifuge tube in a heat block set to 37°C for 30 minutes. After heat shock, embryos were transferred back into E3 water in a petri dish and recovered at 28.5°C. *hsp:dnMAML-GFP* positive embryos were selected based on GFP expression 4 hours after heat shock. For drug treatment, embryos at 18 hpf were treated with 50 μM LY-411575 (Sigma) or DMSO control in E3 fish water until fixation at 30 hpf.

### Notch response analysis

To quantify the total Notch response profiles, *her12:Kaede; vglut2a:dsRed* embryos were imaged at 32 hpf. V3 cells were identified by *vglut2a:dsRed* fluorescence and KA” cells were identified by their characteristic low Notch response and medial localisation. All cells between KA” and V3 cells were defined as LFP progenitors. For each embryo, 10 cells of each type were selected, matching the anterior-posterior positioning of each V3 cell where possible. LFP progenitor cells were always direct neighbours of either V3 or KA” cells. The Kaede fluorescence for each cell was calculated by taking the average fluorescence in a 10-pixel diameter ROI drawn in the centre of the cell. For each embryo, the level of background green fluorescence external to the *her12:Kaede* expression was measured and subtracted from each cell measurement. To analyse the Notch response between 24 hpf and 32 hpf, *her12:Kaede; vglut2a:dsRed* embryos were photoconverted at 24 hpf, imaged directly after, then imaged again at 32 hpf. V3 cells could be identified by an increased level of red fluorescence due to *vglut2a:dsRed* expression compared to neighbouring cells, as well as the lowered Notch response profile visualised in the previous experiment, while KA” and LFP progenitor cells were defined through the same criteria in the previous experiment. The same method as described above was used to quantify Kaede^green^ levels in each cell. To ensure only newly synthesized Kaede was measured, a background value of unconverted Kaede fluorescence was measured and subtracted from the Kaede^green^ measurement. To generate this value, the average Kaede fluorescence of a 200×400 pixel ROI across the converted region of each embryo was taken directly after conversion. The background green fluorescence external to the *her12:Kaede* expression was also measured and subtracted from each cell’s Kaede^green^ measurement. Therefore, the corrected Kaede^green^ signal at 32 hpf = Kaede^green^ measurement at 32 hpf – unconverted Kaede^green^ at 24 hpf – background green signal. In both experiments, each data point displayed is the average of 10 cells. To analyse variance within the groups, the Kruskal-Wallis test was used. To analyse significance between two groups, P values were determined by performing the Mann-Whitney *U* test. All fluorescent imaging was carried out using the Olympus FV1200 confocal microscope with a 20x objective using the Fluoview software. The fluorescent values were measured using Fiji-ImageJ software.

### Quantification of cell numbers

To quantify the number of V3 and KA” cells, embryos were stained with appropriate markers (V3: *sim1*; KA”: *tal2* and *urp1*) and the number of cells from the anterior-most cell to the end of the yolk extension were counted. For quantification of the fluorescent images, cells were counted in the mid-trunk region above the yolk extension.

### Statistical analysis

All the graphs were generated in the GraphPad Prism software. Data were plotted as mean ± SD. Significance between two samples was calculated using Mann-Whitney *U* test. Variance within the groups was calculated using Kruskal-Wallis test. p values: p > 0.05 (not significant, ns); p < 0.05 (*); p < 0.01 (**); p < 0.001 (***); p < 0.0001 (****).

## ACKNOWLEDGEMENTS

We thank the zebrafish community for providing probes and reagents; Jason Berman for *gata2:GFP* fish; Caroline Burns for the *hsp:dnMAML-GFP* plasmid; Sarah Kucenas for the *nkx2.2a:NLS-mCherry* plasmid; Sarah Childs for providing critical input on this project; and members of the Childs and Huang laboratories for discussions.

## COMPETING INTERESTS

The authors declare that no competing interests exist.

## FUNDING

This study was supported by grants to P.H. from the Natural Sciences and Engineering Research Council (NSERC) (RGPIN-2015-06343), Canada Foundation for Innovation John R. Evans Leaders Fund (Project 32920), and Startup Fund from the Alberta Children’s Hospital Research Institute (ACHRI). C.T.J. was supported by Eyes High International Doctoral Scholarship and Alberta Graduate Excellence Scholarship. A.K. was supported by the ACHRI Graduate Scholarship.

---

**Figure S1.**
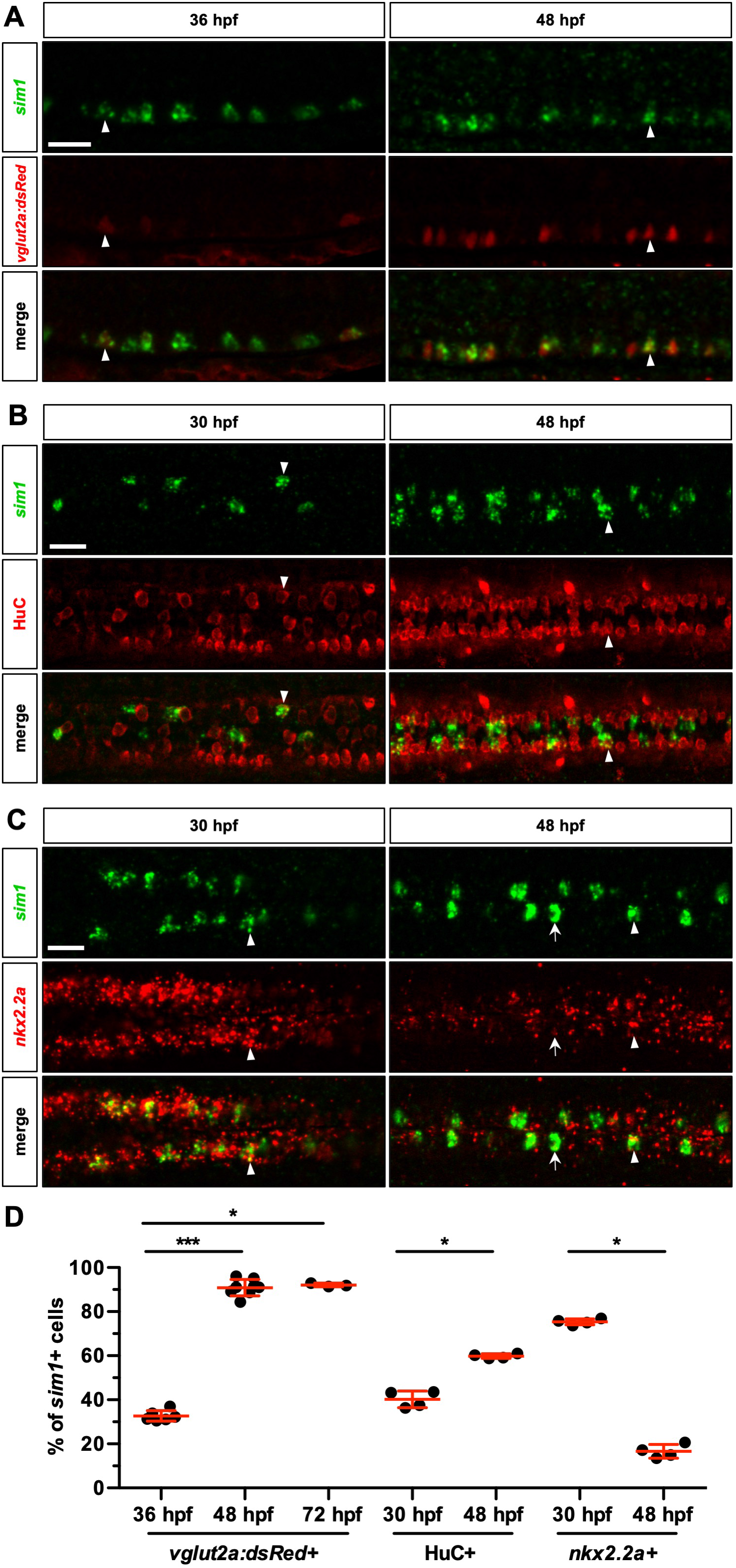
Marker analysis of V3 interneurons in the LFP domain. (A) Whole-mount fluorescent in situ hybridisation for *sim1* was performed alongside immunohistochemistry for dsRed on *vglut2a:dsRed* embryos at either 36 hpf or 48 hpf. Fixed embryos were imaged from the lateral view. Arrowheads highlight cells that are positive for both *sim1* and dsRed. (B) Whole-mount fluorescent in situ hybridisation for *sim1* was performed alongside immunohistochemistry for HuC on wild type embryos at either 30 hpf or 48 hpf. Fixed embryos were imaged from the dorsal view. Arrowheads highlight cells that are positive for both *sim1* and HuC. (C) Whole-mount double fluorescent in situ hybridisation for *sim1* and *nkx2.2a* was performed on wild type embryos at either 30 hpf or 48 hpf. Fixed embryos were imaged from the dorsal view. Arrowheads highlight cells that are positive for both *sim1* and *nkx2.2a*. Arrows highlight *sim1^+^* cells that are negative for *nkx2.2a*. (D) The number of double positive cells are displayed as a percentage of the total *sim1* positive cell population in each condition. Each data point represents the percentage of double positive cells in one embryo. Data are plotted with mean ± SD. Statistics: Mann-Whitney *U* test. Asterisks representation: p-value<0.05 (*), p-value<0.001 (***). *n* = 3-8 embryos per condition. Scale bars: 20 μm.

**Figure S2.**
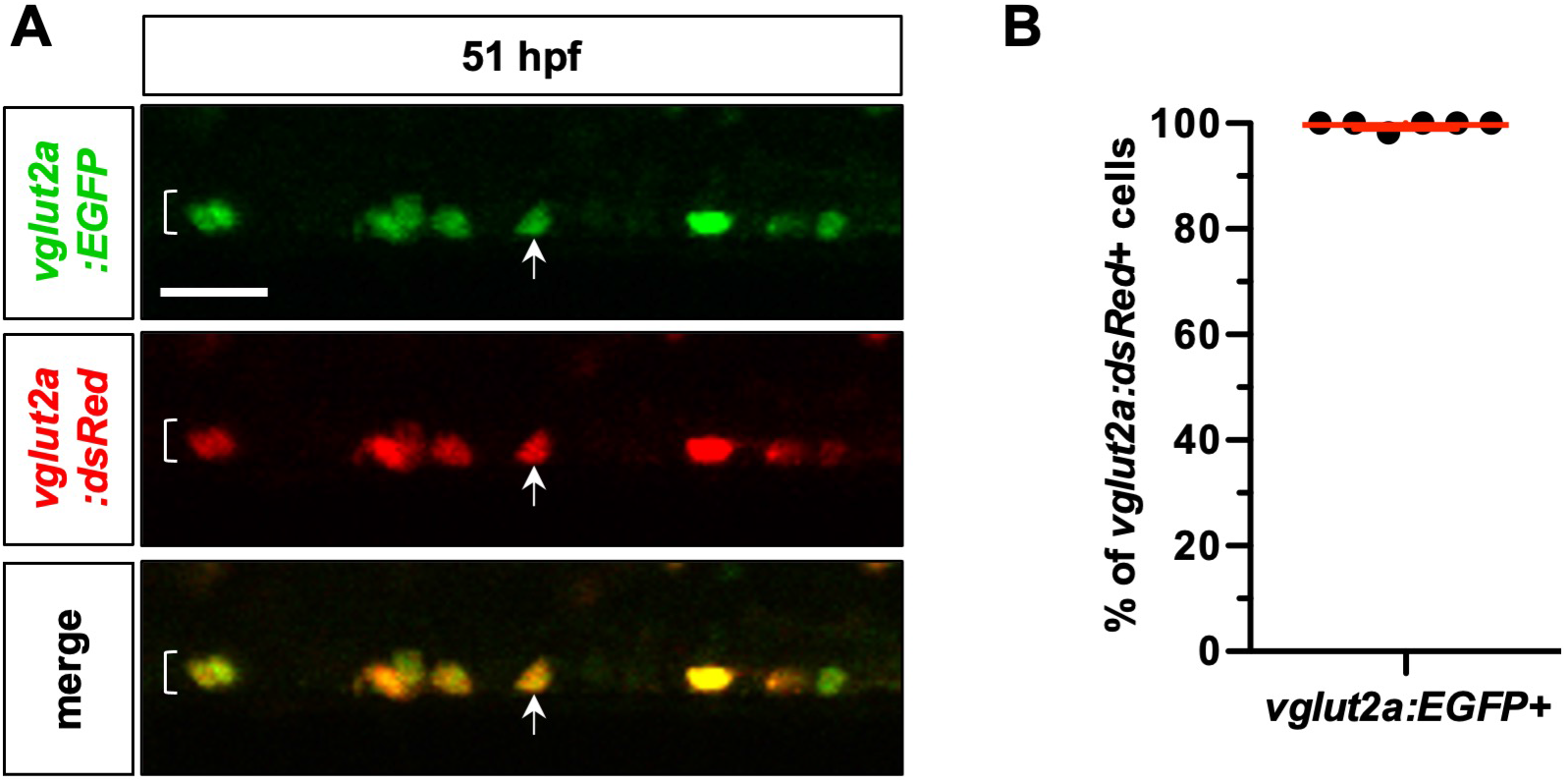
The *vglut2a:EGFP* and *vglut2a:dsRed* reporters label an identical cell population in the LFP domain. (A) *vglut2a:EGFP; vglut2a:dsRed* embryos were imaged at 51 hpf. Images are taken from the lateral view. Bracket denotes the lateral floor plate domain. Arrows denote a double positive cell. (B) The number of *vglut2a:EGFP* and *vglut2a:dsRed* double positive cells displayed as a percentage of the total *vglut2a:dsRed* positive cell population. Each data point represents the percentage of double positive cells in one embryo. The image shown in (A) is an example of one of these embryos. Data are plotted with mean ± SD. *n* = 6 embryos. Scale bar: 20 μm.

**Figure S3.**
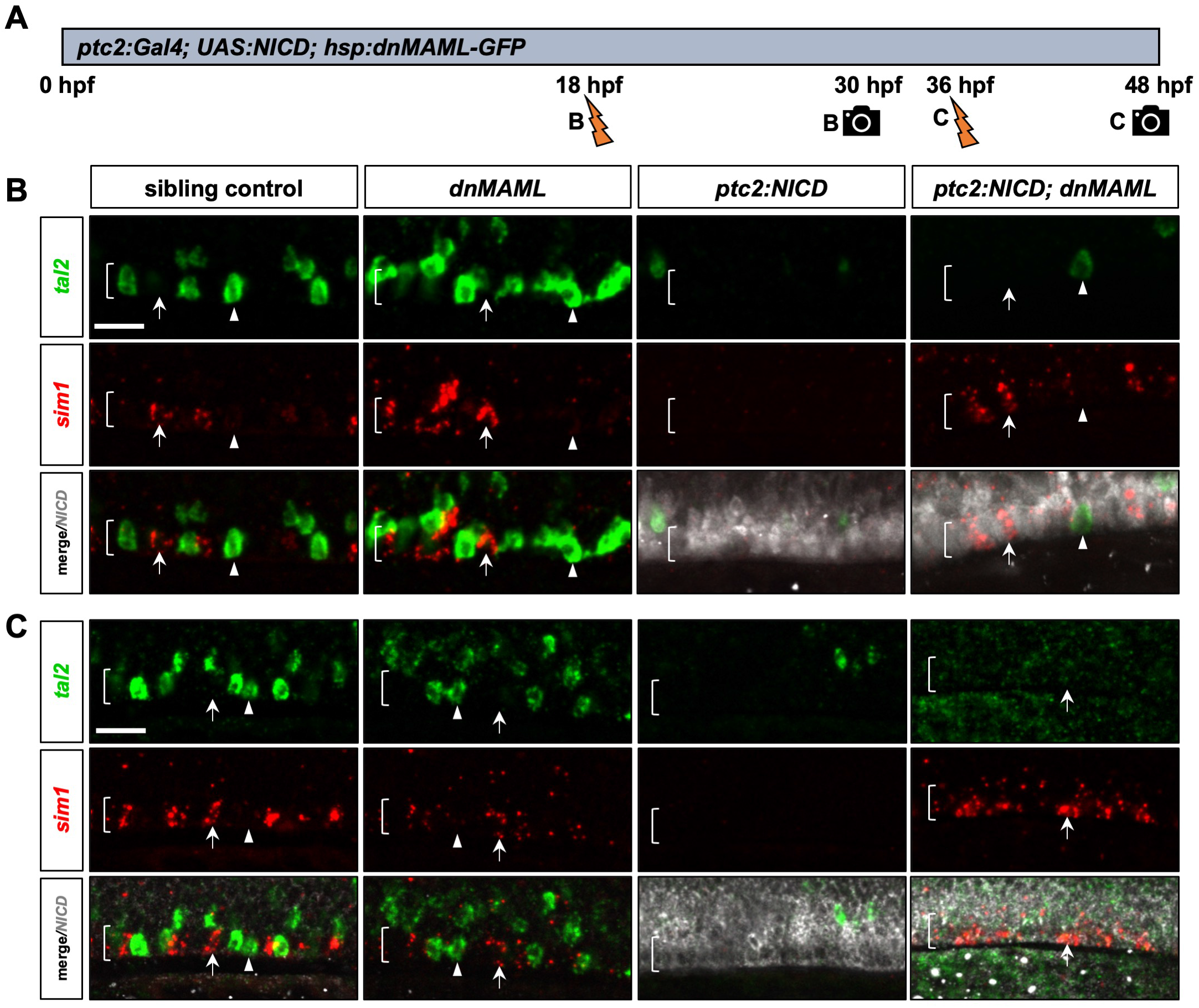
The duration of Notch signalling mediates the different waves of differentiation. (A) Schematic representation of the experimental design for (B) and (C). (B, C) *hsp:dnMAML-GFP* (shown as *dnMAML*), *ptc2:Gal4; UAS:NICD* (shown as *ptc2:NICD*), *ptc2:NICD; hsp:dnMAML-GFP* and their wild type sibling embryos were heat shocked at either 18 hpf and fixed at 30 hpf (B) or heat shocked at 36 hpf and then fixed at 48 hpf (C). Whole-mount double fluorescent in situ hybridisation for *tal2* and *sim1* was performed alongside immunohistochemistry for Myc to label Myc tagged NICD expression. Fixed embryos were imaged from the lateral view. Arrowheads highlight *tal2^+^* KA” cells. Arrows highlight *sim1^+^* V3 cells. Brackets denote the extent of the LFP domain. *n* = 4 embryos per condition. Scale bars: 20 μm.

**Figure S4.**
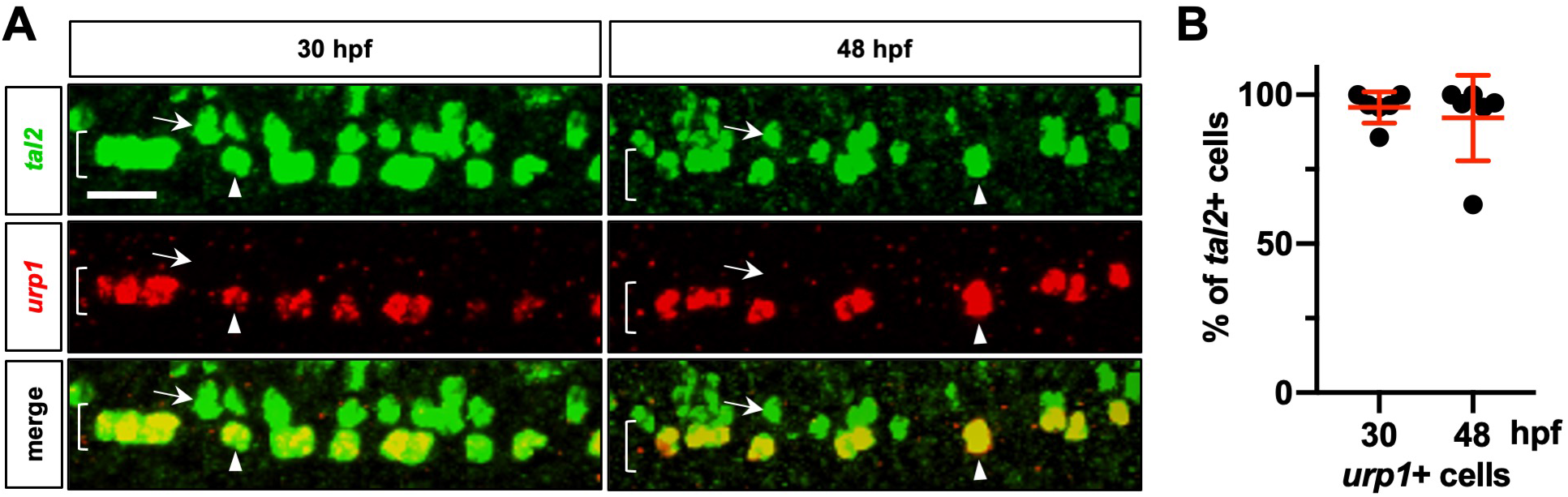
KA” cells are specifically labelled by *urp1* expression. (A) Whole-mount double fluorescent in situ hybridisation for *tal2* and *urp1* was performed at 30 hpf and 48 hpf. Fixed embryos were imaged from the lateral view. Arrowheads highlight *tal2^+^urp1^-^* KA” cells. Arrows highlight *tal2+urp1^+^* KA’ cells. Brackets denote the extent of the LFP domain. *n* = 6 embryos per condition. (B) The number of *tal2^+^urp1^+^* cells are displayed as a percentage of the total *tal2^+^* cell population at each time point. Each data point represents the percentage of double positive cells in one embryo. The images shown in (A) are examples of these embryos. Data are plotted with mean ± SD. Scale bars: 20 μm.

## Notes

### Competing Interest Statement

The authors have declared no competing interest.

## REFERENCES

Agirman, G., Broix, L. and Nguyen, L. (2017). Cerebral cortex development: an outside-in perspective. FEBS Lett. 591, 3978–3992.

Alaynick, W. A., Jessell, T. M. and Pfaff, S. L. (2011). SnapShot: Spinal Cord Development. Cell 146, 178–178.e1.

Andrews, M. G., Kong, J., Novitch, B. G. and Butler, S. J. (2019). New perspectives on the mechanisms establishing the dorsal-ventral axis of the spinal cord. Curr. Top. Dev. Biol. 132, 417–450.

Appel, B., Givan, L. A. and Eisen, J. S. (2001). Delta-Notch signaling and lateral inhibition in zebrafish spinal cord development. BMC Dev. Biol. 1, 1–11.

Artavanis-Tsakonas, S. and Simpson, P. (1991). Choosing a cell fate: a view from the Notch locus. Trends Genet. 7, 403–408.

Bae, Y. K., Kani, S., Shimizu, T., Tanabe, K., Nojima, H., Kimura, Y., Higashijima, S. ichi and Hibi, M. (2009). Anatomy of zebrafish cerebellum and screen for mutations affecting its development. Dev. Biol. 330, 406–426.

Batista, M. F., Jacobstein, J. and Lewis, K. E. (2008). Zebrafish V2 cells develop into excitatory CiD and Notch signalling dependent inhibitory VeLD interneurons. Dev. Biol. 322, 263–275.

Briscoe, J. and Small, S. (2015). Morphogen rules: Design principles of gradient-mediated embryo patterning. Dev. 142, 3996–4009.

Briscoe, J., Pierani, A., Jessell, T. M. and Ericson, J. (2000). A homeodomain protein code specifies progenitor cell identity and neuronal fate in the ventral neural tube. Cell 101, 435–445.

Bussmann, J. and Schulte-Merker, S. (2011). Rapid BAC selection for tol2-mediated transgenesis in zebrafish. Development 138, 4327–4332.

Cohen, M., Briscoe, J. and Blassberg, R. (2013). Morphogen interpretation: the transcriptional logic of neural tube patterning. Curr. Opin. Genet. Dev. 23, 423–428.

Davison, J. M., Akitake, C. M., Goll, M. G., Rhee, J. M., Gosse, N., Baier, H., Halpern, M. E., Leach, S. D. and Parsons, M. J. (2007). Transactivation from Gal4-VP16 transgenic insertions for tissue-specific cell labeling and ablation in zebrafish. Dev. Biol. 304, 811–824.

Del Barrio, M. G., Taveira-Marques, R., Muroyama, Y., Yuk, D.-I., Li, S., Wines-Samuelson, M., Shen, J., Smith, H. K., Xiang, M., Rowitch, D., et al. (2007). A regulatory network involving Foxn4, Mash1 and delta-like 4/Notch1 generates V2a and V2b spinal interneurons from a common progenitor pool. Development 134, 3427–3436.

Dessaud, E., Yang, L. L., Hill, K., Cox, B., Ulloa, F., Ribeiro, A., Mynett, A., Novitch, B. G. and Briscoe, J. (2007). Interpretation of the sonic hedgehog morphogen gradient by a temporal adaptation mechanism. Nature 450, 717–720.

Dessaud, E., McMahon, A. P. and Briscoe, J. (2008). Pattern formation in the vertebrate neural tube: A sonic hedgehog morphogen-regulated transcriptional network. Development 135, 2489–2503.

Fauq, A. H., Simpson, K., Maharvi, G. M., Golde, T. and Das, P. (2007). A multigram chemical synthesis of the γ-secretase inhibitor LY411575 and its diastereoisomers. Bioorganic Med. Chem. Lett. 17, 6392–6395.

Formosa-Jordan, P., Ibañes, M., Ares, S. and Frade, J.-M. (2013). Lateral inhibition and neurogenesis: novel aspects in motion. Int. J. Dev. Biol. 57, 341–350.

Guruharsha, K. G., Kankel, M. W. and Artavanis-Tsakonas, S. (2012). The Notch signalling system: recent insights into the complexity of a conserved pathway. Nat. Rev. Genet. 13, 654–666.

Huang, P., Xiong, F., Megason, S. G. and Schier, A. F. (2012). Attenuation of Notch and Hedgehog signaling is required for fate specification in the spinal cord. PLoS Genet. 8, e1002762.

Jacobs, C. T. and Huang, P. (2019). Notch signalling maintains hedgehog responsiveness via a gli-dependent mechanism during spinal cord patterning in zebrafish. Elife 8, 1–24.

Jessell, T. M. (2000). Neuronal specification in the spinal cord: Inductive signals and transcriptional codes. Nat. Rev. Genet. 1, 20–29.

Kageyama, R., Ohtsuka, T., Shimojo, H. and Imayoshi, I. (2008). Dynamic Notch signaling in neural progenitor cells and a revised view of lateral inhibition. Nat. Neurosci. 11, 1247–1251.

Kimura, Y., Satou, C. and Higashijima, S. (2008). V2a and V2b neurons are generated by the final divisions of pair-producing progenitors in the zebrafish spinal cord. Development 135, 3001–3005.

Kong, J. H., Yang, L., Dessaud, E., Chuang, K., Moore, D. M., Rohatgi, R., Briscoe, J. and Novitch, B. G. (2015). Notch Activity Modulates the Responsiveness of Neural Progenitors to Sonic Hedgehog Signaling. Dev. Cell 33, 373–87.

Kopan, R. and Ilagan, M. X. G. (2009). The Canonical Notch Signaling Pathway: Unfolding the Activation Mechanism. Cell 137, 216–233.

Kucenas, S., Takada, N., Park, H. C., Woodruff, E., Broadie, K. and Appel, B. (2008). CNS-derived glia ensheath peripheral nerves and mediate motor root development. Nat. Neurosci. 11, 143–151.

Le Dréau, G. and Martí, E. (2012). Dorsal-ventral patterning of the neural tube: A tale of three signals. Dev. Neurobiol. 72, 1471–1481.

Lewis, K. E. and Eisen, J. S. (2003). From cells to circuits: development of the zebrafish spinal cord. Prog. Neurobiol. 69, 419–449.

Liu, Y., Pathak, N., Kramer-Zucker, A. and Drummond, I. A. (2007). Notch signaling controls the differentiation of transporting epithelia and multiciliated cells in the zebrafish pronephros. Development 134, 1111–1122.

Louvi, A. and Artavanis-Tsakonas, S. (2006). Notch signalling in vertebrate neural development. Nat Rev Neurosci 7, 93–102.

Marklund, U., Hansson, E. M., Sundstrom, E., de Angelis, M. H., Przemeck, G. K. H., Lendahl, U., Muhr, J. and Ericson, J. (2010). Domain-specific control of neurogenesis achieved through patterned regulation of Notch ligand expression. Development 137, 437–445.

Miyasaka, N., Morimoto, K., Tsubokawa, T., Higashijima, S. I., Okamoto, H. and Yoshihara, Y. (2009). From the olfactory bulb to higher brain Centers: Genetic visualization of secondary olfactory pathways in zebrafish. J. Neurosci. 29, 4756–4767.

Noctor, S. C., Martinez-Cerdeño, V., Ivic, L. and Kriegstein, A. R. (2004). Cortical neurons arise in symmetric and asymmetric division zones and migrate through specific phases. Nat. Neurosci. 7, 136–144.

Odenthal, J., Van Eeden, F. J. M., Haffter, P., Ingham, P. W. and Nüsslein-Volhard, C. (2000). Two distinct cell populations in the floor plate of the Zebrafish are induced by different pathways. Dev. Biol. 219, 350–363.

Okigawa, S., Mizoguchi, T., Okano, M., Tanaka, H., Isoda, M., Jiang, Y. J., Suster, M., Higashijima, S. ichi, Kawakami, K. and Itoh, M. (2014). Different combinations of Notch ligands and receptors regulate V2 interneuron progenitor proliferation and V2a/V2b cell fate determination. Dev. Biol. 391, 196–206.

Park, H. C. and Appel, B. (2003). Delta-Notch signaling regulates oligodendrocyte specification. Development 130, 3747–3755.

Peng, C.-Y., Yajima, H., Burns, C. E., Zon, L. I., Sisodia, S. S., Pfaff, S. L. and Sharma, K. (2007). Notch and MAML Signaling Drives Scl-Dependent Interneuron Diversity in the Spinal Cord. Neuron 53, 813–827.

Petracca, Y. L., Sartoretti, M. M., Di Bella, D. J., Marin-Burgin, A., Carcagno, A. L., Schinder, A. F. and Lanuza, G. M. (2016). The late and dual origin of cerebrospinal fluid-contacting neurons in the mouse spinal cord. Development 143, 880–891.

Pierfelice, T., Alberi, L. and Gaiano, N. (2011). Notch in the Vertebrate Nervous System: An Old Dog with New Tricks. Neuron 69, 840–855.

Quan, F. B., Dubessy, C., Galant, S., Kenigfest, N. B., Djenoune, L., Leprince, J., Wyart, C., Lihrmann, I. and Tostivint, H. (2015). Comparative Distribution and In Vitro Activities of the Urotensin II-Related Peptides URP1 and URP2 in Zebrafish: Evidence for Their Colocalization in Spinal Cerebrospinal Fluid-Contacting Neurons. PLoS One 10,.

Sadowski, I., Ma, J., Triezenberg, S. and Ptashne, M. (1988). GAL4-VP16 is an unusually potent transcriptional activator. Nature 335, 563–564.

Sato, M., Yasugi, T., Minami, Y., Miura, T. and Nagayama, M. (2016). Notch-mediated lateral inhibition regulates proneural wave propagation when combined with EGF-mediated reaction diffusion. Proc. Natl. Acad. Sci. U. S. A. 113, E5153–E5162.

Schäfer, M., Kinzel, D., Neuner, C., Schartl, M., Volff, J. N. and Winkler, C. (2005). Hedgehog and retinoid signalling confines nkx2.2b expression to the lateral floor plate of the zebrafish trunk. Mech. Dev. 122, 43–56.

Schäfer, M., Kinzel, D. and Winkler, C. (2007). Discontinuous organization and specification of the lateral floor plate in zebrafish. Dev. Biol. 301, 117–129.

Scheer, N. and Campos-Ortega, J. A. (1999). Use of the Gal4-UAS technique for targeted gene expression in the zebrafish. Mech. Dev. 80, 153–158.

Shin, J., Poling, J., Park, H.-C. and Appel, B. (2007). Notch signaling regulates neural precursor allocation and binary neuronal fate decisions in zebrafish. Development 134, 1911–1920.

Stasiulewicz, M., Gray, S., Mastromina, I., Silva, J. C., Bjorklund, M., Seymour, P. a, Booth, D., Thompson, C., Green, R., Hall, E. a, et al. (2015). A conserved role for Notch in priming the cellular response to Shh through ciliary localisation of the key Shh transducer, Smoothened. Development 2291–2303.

Traver, D., Paw, B. H., Poss, K. D., Penberthy, W. T., Lin, S. and Zon, L. I. (2003). Transplantation and in vivo imaging of multilineage engraftment in zebrafish bloodless mutants. Nat. Immunol. 4, 1238–1246.

Ulloa, F. and Martí, E. (2010). Wnt won the war: Antagonistic role of Wnt over Shh controls dorso-ventral patterning of the vertebrate neural tube. Dev. Dyn. 239, 69–76.

Vize, P. D., McCoy, K. E. and Zhou, X. (2009). Multichannel wholemount fluorescent and fluorescent/chromogenic in situ hybridization in Xenopus embryos. Nat. Protoc. 4, 975–83.

Yang, X., Tomita, T., Wines-Samuelson, M., Beglopoulos, V., Tansey, M. G., Kopan, R. and Shen, J. (2006). Notch1 Signaling Influences V2 Interneuron and Motor Neuron Development in the Spinal Cord. Dev. Neurosci. 28, 102–117.

Yang, L., Rastegar, S. and Strähle, U. (2010). Regulatory interactions specifying Kolmer–Agduhr interneurons. Development 137, 2713–2722.

Yang, L., Wang, F. and Strähle, U. (2020). The Genetic Programs Specifying Kolmer-Agduhr Interneurons. Front. Neurosci. 14, 1064.

Yeo, S.-Y. and Chitnis, A. B. (2007). Jagged-mediated Notch signaling maintains proliferating neural progenitors and regulates cell diversity in the ventral spinal cord. Proc. Natl. Acad. Sci. U. S. A. 104, 5913–8.

Yoon, K. and Gaiano, N. (2005). Notch signaling in the mammalian central nervous system: Insights from mouse mutants. Nat. Neurosci. 8, 709–715.

Zhao, L., Borikova, A. L., Ben-Yair, R., Guner-Ataman, B., MacRae, C. A., Lee, R. T., Geoffrey Burns, C. and Burns, C. E. (2014). Notch signaling regulates cardiomyocyte proliferation during zebrafish heart regeneration. Proc. Natl. Acad. Sci. U. S. A. 111, 1403–1408.

Zhu, Y., Crowley, S. C., Latimer, A. J., Lewis, G. M., Nash, R. and Kucenas, S. (2019). Migratory Neural Crest Cells Phagocytose Dead Cells in the Developing Nervous System. Cell 179, 74–89.e10.

